# Versatile and efficient non-viral integration of large transgenes in human T cells via CRISPR knock-in and engineered integrases

**DOI:** 10.1101/2025.09.10.675267

**Authors:** Isabell Kassing, Jonas Kath, Ana-Maria Nitulescu, Viktor Glaser, Laura M. Hartmann, Yaolin Pu, Luis Huth, Roberts Kārkliņš, Shona Shaji, Alessa R. Ringel, Marie Pouzolles, Maik Stein, Daniel M. Ibrahim, Dimitrios L. Wagner

**Affiliations:** Berlin Center for Advanced Therapies, Charité-Universitätsmedizin Berlin, Corporate Member of Freie Universität Berlin, Humboldt-Universität zu Berlin, and Berlin Institute of Health, Berlin, Germany; Berlin Institute of Health Center for Regenerative Therapies, Berlin Institute of Health at Charité-Universitätsmedizin Berlin, Berlin, Germany; Center for Cell and Gene Therapy, Baylor College of Medicine, Houston, TX; Institute of Medical Immunology, Charité-Universitätsmedizin Berlin, Corporate Member of Freie Universität Berlin, Humboldt-Universität zu Berlin, and Berlin Institute of Health, Berlin, Germany; Max-Planck Institute for Molecular Genetics, Berlin, Germany; Department of Molecular and Cellular Biology, Baylor College of Medicine, Houston, TX; Dan L. Duncan Comprehensive Cancer Center, Baylor College of Medicine, Houston, TX

## Abstract

Current gene transfer methods often lack the precision, versatility, or efficiency when integrating large transgenes, limiting the ability to engineer therapeutic T-cells with more complex payloads. Here, we report ‘one-pot’ PASTA (Programmable and Site-specific Transgene Addition), a non-viral genome engineering strategy for large gene insertion that combines CRISPR-Cas-mediated homology-directed repair (HDR) and site-specific recombination via serine integrases. Using ‘one-pot’ PASTA with the Bxb1 integrase, we demonstrate efficient integration of transgenes at multiple genomic loci relevant for T-cell engineering (e.g., *TRAC, B2M, CD3E, CD3Z, GAPDH*). For constructs > 8 kb, ‘one-pot’ PASTA outperforms conventional HDR by 19-fold on average and prime-editing-assisted site-specific integrase gene editing (PASSIGE) by 5-fold. This enables the delivery of multi-cistronic cargo to generate dual-antigen targeting CAR T-cells with a safety-switch that overcome antigen escape in lymphoma models. Finally, ‘one-pot’ PASTA can be further optimized with improved integrase enzymes, such as engineered variants of Pa01 or Bxb1, and plasmids with minimized backbones. In summary, ‘one-pot’ PASTA represents a versatile and scalable platform for precise, non-viral gene insertion in T-cells.

## Introduction

Adoptive T-cell therapy, particularly using chimeric antigen receptor (CAR) T cells, has revolutionized the treatment of B cell malignancies and autoimmune diseases^1,2^. However, for many disorders, such as solid cancers, CAR T cells are not yet effective, or patients relapse after initial remission^3^. Antigen escape prevents long-term cures with single-antigen targeted CAR T cell therapies^4^. Patient-derived T cells often lose function within the tumor microenvironment due to insufficient cytokine support and immunosuppressive signals, leading to exhaustion and reduced therapeutic efficacy^3,5^. CAR T cells can also cause significant side-effects through neurotoxicity or on-target off-tumor toxicity, limiting the doses and utility^3,6,7^. To overcome these challenges, many strategies have been proposed, such as co-expression of multiple antigen receptors, cytokines, and safety switches or spatiotemporal regulation of CARs, enhancing the overall therapeutic efficacy and safety of CAR T cell therapies^6^. However, these approaches require gene transfer of large and complex genetic payloads which remains a major challenge, particularly in therapeutically relevant cell types.

Various methods for gene transfer in T cells have been explored, each with its own set of advantages and limitations. Non-targeted integration methods, such as lentiviral (LV) and retroviral transduction, as well as transposase systems, are commonly used but carry risks of insertional mutagenesis and unpredictable gene expression^8–11^. On the other hand, targeted integration methods, including homology-directed repair (HDR) using non-viral vectors (pDNA^12,13^, linear dsDNA^14–16^, linear ssDNA^14,17,18^, circular ssDNA^19^) and adeno-associated virus (AAV) vectors^20–22^, offer more precise gene insertion but are often limited by cargo size and low efficiency for transgene sizes exceeding the AAV packaging capacity (e.g. 5kb, without homology arms).

Recently, techniques like PASSIGE^23^ and PASTE^24^ have addressed some of these limitations by combining prime editing (PE) with large serine recombinases (LSRs or serine integrases), such as Bxb1. LSRs are bacteriophage-derived DNA integrases that mediate site-specific recombination between short DNA sequences called attachment sites, typically attP (in phages) and attB (in bacteria)^25^. This unidirectional reaction joins halves of attP and attB to form new hybrid sites, attL (left) and attR (right), flanking the integrated DNA. To harness this mechanism for mammalian genome engineering, a synthetic landing pad (attG – corresponding to either attB or attP) can be introduced at a precise genomic locus, enabling site-specific integration of a DNA donor vector carrying the matching attachment site (attV – attP or attB, respectively). While attB and attP sites do not naturally occur in the human genome, similar sequences, known as pseudosites, can be present and may lead to potential off-target recombination events^26,27^. In PASSIGE or PASTE, PE is used to insert an attG site/landing pad with two complementary prime editors (twin-PE), since insertions over 40 bp are inefficient with a single prime editor^23,24^. The corresponding LSR is provided as part of a fusion protein (PASTE) or by co-transfection of a plasmid (PASSIGE) and then catalyzes the landing pad-directed integration of a DNA cargo, resulting in site-specific genomic integration. However, while PE can be efficient for landing pad insertions in primary cells^23,28,29^, its efficient adaption to new target sites remains technically challenging and requires highly optimized reagents, which limits its versatility. For instance, efficient PE in primary cells requires long, chemically protected guide RNAs which push the limits of current solid state synthesis methods^30^.

Here, we set out to establish a versatile nuclease-mediated approach, which we term ‘one-pot’ PASTA (Programmable and Site-specific Transgene Addition), combining LSRs with conventional Cas-mediated knock-in of landing pads in a single transfection. In contrast to PE-alternatives, HDR is easily adaptable to novel sites, relies on broadly available CRISPR-Cas reagents, and is highly efficient in primary human T cells – particularly for small transgenes, such as landing pads. Leveraging this unique combination of established methods, ‘one-pot’ PASTA demonstrates high efficiency across different sites in primary human T cells, with greater integration efficacies compared to conventional non-viral HDR for transgenes exceeding 6 kb and improved recombination rates over PASSIGE. We demonstrate that the increased cargo capacity allows optimization of CAR T cell therapy in a lymphoma model of antigen escape, highlighting its utility in a clinically relevant context. Finally, to further enhance ‘one-pot’ PASTA, we screened multiple integrases for their utility in primary human T cells and generated an enhanced Pa01 LSR variant, which outperforms its wildtype version and the original Bxb1 LSR. In summary, ‘one-pot’ PASTA represents a scalable platform for non-viral, site-specific gene insertion in T cells, offering a versatile and efficient alternative to prime editing-based approaches.

## Results

### ‘One-pot’ PASTA achieves consistently high CAR integration at various genomic loci

To establish a versatile genome engineering approach, we developed ‘one-pot’ PASTA, a single electroporation platform enabling site-specific integration of large circular DNA payloads in primary human T cells (**Fig. 1A**). The system relies on the delivery of four components: (**1**) Cas-crRNA ribonucleoprotein (RNP), (**2**) a linear double-stranded DNA template designed for HDR-mediated insertion of a 38 bp attachment site (“landing pad”) for the LSR Bxb1, (**3**) Bxb1 mRNA, and (**4**) a circular plasmid carrying the transgene of interest and the corresponding attachment site. The initially described attB was selected as genomic attachment site (attG), whereas attP served as attachment site on the circular DNA vector (attV). The editing occurs in two steps: HDR-mediated insertion of the landing pad, followed by Bxb1-mediated site-specific integration of the plasmid (**Fig. 1A**). To validate the platform, we integrated a CD19-CAR lacking an exogenous promotor (Plasmid #1) into five transcriptionally active loci that are relevant to T cell engineering (*TRAC* exon 1 [Ref.^16,31^], *CD3E* exon 3 [Ref.^32^], *CD3Z* exon 2 [Ref.^33^], *B2M* exon 1 [Ref. ^34^], and *B2M* exon 2 [Ref.^34^]) with either Cas9- or Cas12a-RNPs. Editing outcomes were assessed by flow cytometry four days after electroporation (**Fig. 1B-D**). Control conditions lacking one or more components (mock-electroporation, RNP only, no plasmid, plasmid only) showed no detectable CAR expression (**Fig. 1B**). In contrast, stable CAR integration was observed only when all four components were delivered, with mean CAR integration rates ranging from 29% (*CD3Z*) to 44% (*B2M* exon 1). CAR surface expression levels differed by integration site, reflecting the activity of the respective endogenous promoters, with the lowest expression observed at *CD3Z* and the highest at *B2M* exon 2. Mean survival across all five targeted loci exceeded 75% relative to mock-electroporated controls (**Fig. 1D**). To optimize editing conditions, we systematically compared two electroporation programs (EH-115 vs. EO-115) and post-electroporation treatments (± HDR-enhancing DNA-PK inhibitor AZD7648 and ART558), intended to favor precise HDR over non-homologous end-joining^35,36^ (**Suppl. Fig. 1A, B**). Electroporation program EH-115 and AZD7648/ART558 treatment yielded the highest integration rates, outweighing the slightly improved viability observed with EO-115. Ultimately, across all five integration sites, the resulting CAR T cells showed strong CAR-dependent cytotoxicity against CD19⁺ Nalm-6 target cells in co-culture assays, confirming functional potency *in vitro* (**Fig. 1E**).

**Fig. 1.**
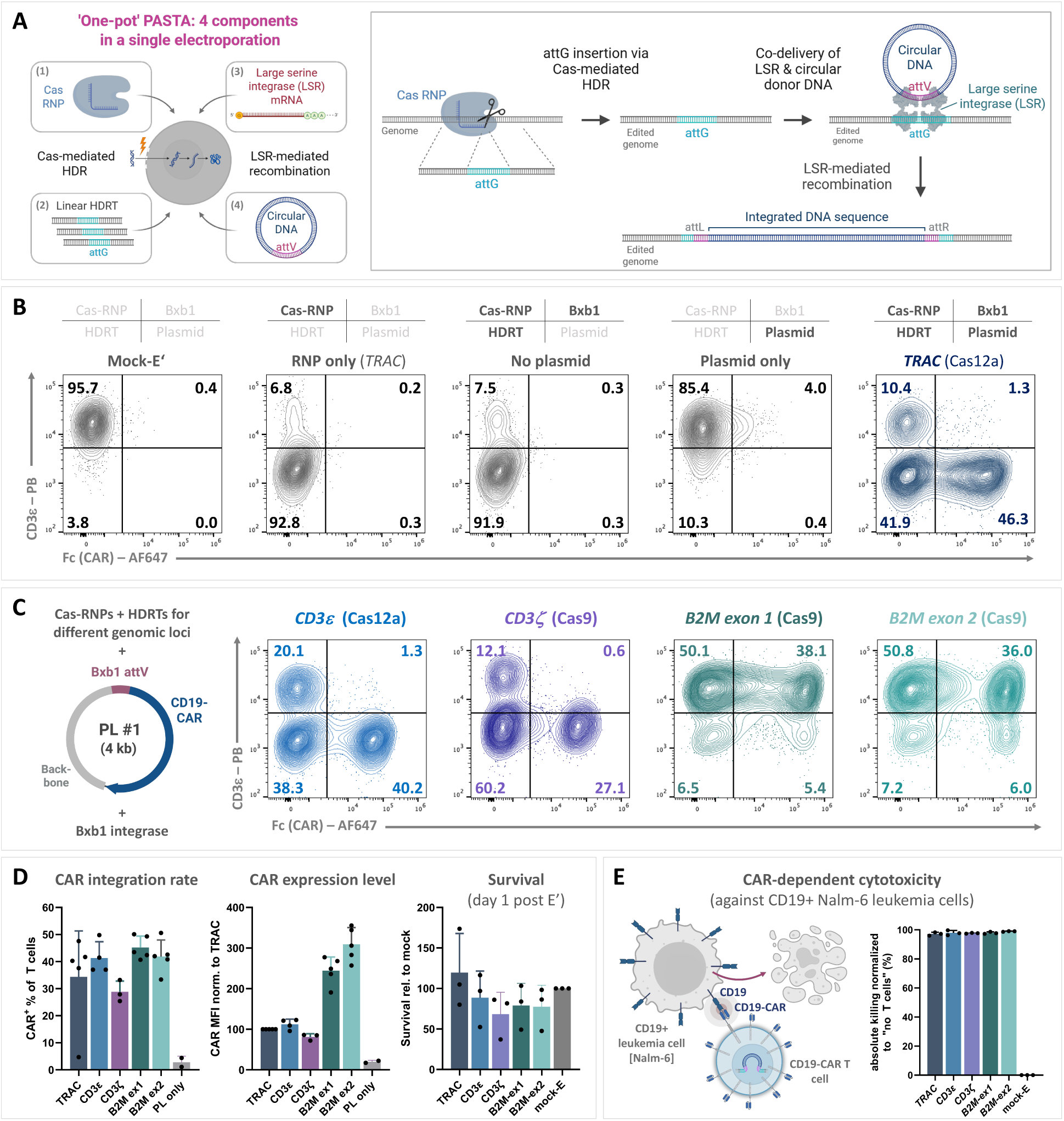
One-pot PASTA – an efficient and versatile system for site-specific integration of transgenes. (**A**) Schematic overview of the ‘one-pot’ PASTA system consisting of four components. RNP: Ribonucleoprotein; HDRT: homology-directed repair template; LSR: large serine integrase; attG: attachment site in genome; attV: attachment site on vector; attL/R: recombined attachment site left/right. (**B**) Bxb1-mediated integration of a CD19-CAR into the *TRAC* locus of primary human T cells. Representative flow cytometry plots show different editing outcomes when omitting individual components of the PASTA system. Flow cytometry analysis was performed on day 4 after electroporation. (**C**) Representation of plasmid (PL) #1 for integration of a CD19-directed CAR. Representative flow cytometry plots demonstrate Bxb1-mediated CD19-CAR integration with landing pad insertion at alternative T cell-relevant genomic loci (*CD3ε, CD3ζ, B2M exon1, B2M exon2*) by adaption of Cas-RNP (Cas9 or Cas12a) and HDR template respectively. Flow cytometry analysis was performed on day 4 post electroporation. (**D**) CD19-CAR integration at different genomic loci in primary human T cells. Summarized data show integration rates, CAR expression levels (measured as median fluorescence intensity, MFI) and survival of cells 24h after electroporation, all determined by flow cytometry (n = 3-5 healthy donors). (**E**) CAR-dependent cytotoxicity against CD19^+^ Nalm-6 leukemia cells assessed by flow cytometry after 24h of co-culture in (n = 3 healthy donors).

### ‘One-pot’ PASTA enables efficient integration of large and complex transgene cassettes

To assess the capacity of ‘one-pot’ PASTA for delivery of increasingly complex genetic payloads, we tested three transgene cassettes of ascending size: a monocistronic construct encoding a CD19-CAR (Plasmid (PL) #1), a bicistronic cassette encoding CD19-CAR and a truncated EGFR (tEGFR) lacking intracellular signaling domains (PL #2), and a tricistronic cassette encoding CD19-CAR, tEGFR, and a CD20-CAR (PL #3) (**Fig. 2A**). Integration plasmid sizes, including the plasmid backbone, were around 4 kb, 7 kb and 8.5 kb, respectively. While the CD19-CAR lacked a promoter, both the bi- and tricistronic constructs included an EF1α promoter upstream of the tEGFR module. Flow cytometry confirmed expression of all expected markers: CD19-CAR was detected in all three configurations, tEGFR in the bi- and tricistronic settings, and CD20-CAR exclusively in the tricistronic construct (**Fig. 2B**). Notably, increasing plasmid size did not negatively impact the integration efficiency of ‘one-pot’ PASTA, measured via CD19-CAR expression from the *TRAC* locus as a stringent readout of stable genomic integration (**Fig. 2C**.) In contrast, conventional HDR strategies relying on full-length transgene insertion exhibited a marked drop in efficiency with larger constructs (**Fig. 2D**). For the 6.5 kb transgene cassette (PL #3), this resulted in a significant 19-fold average increase in integration efficiency with ‘one-pot’ PASTA compared to conventional HDR (**Fig. 2E**). Assessment of post-editing cell viability did not reveal toxicity associated with larger plasmids during ‘one-pot’ PASTA, whereas conventional HDR showed a decline in cell viability with increasing template size (**Suppl. Fig. 2**). These findings demonstrate that ‘one-pot’ PASTA enables efficient and scalable integration of large, multigenic payloads.

**Fig. 2.**
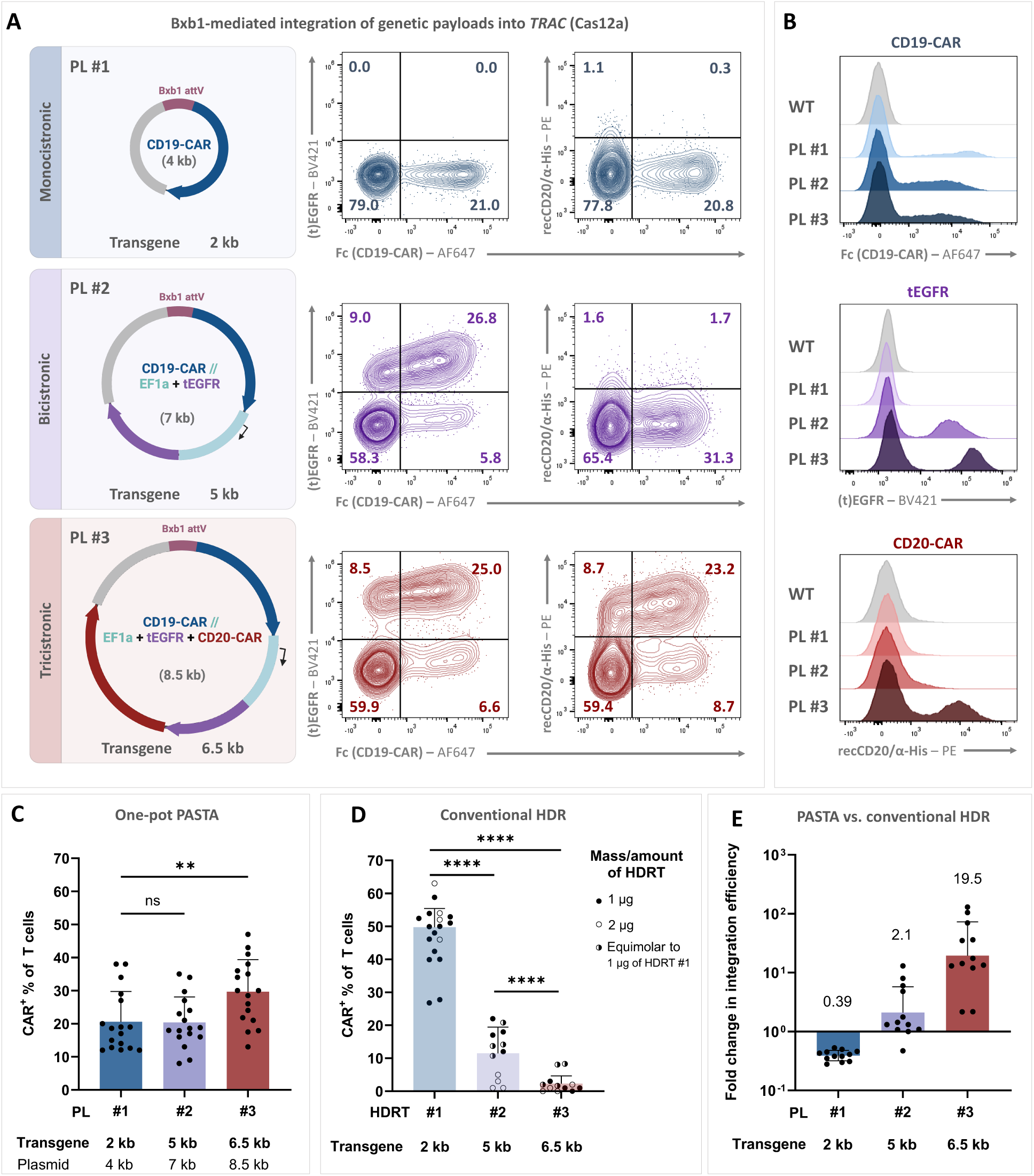
PASTA outperforms conventional HDR for integration of large transgenes. (**A**) Representative flow cytometry plots for integration of plasmids with increasing genetic payloads into *TRAC*. Plasmid 1 (PL #1) encodes a CD19-directed CAR, plasmid 2 (PL #2) includes a truncated epidermal growth factor (tEGFR) driven by an exogenous EF1a-promoter downstream of the CD19-CAR, and plasmid 3 (PL #3) carries an additional CD20-CAR downstream of tEGFR. (**B**) Expression of individual transgenes in CD3-T cells. (**C**) Integration of plasmids PL #1-3 via ‘one-pot’ PASTA. All plasmids were transfected in equimolar amounts of 0.6 nmol. (n = 17 healthy donors; Friedman test of matched data followed by Dunńs multiple comparisons test). (**D**) Insertion of transgenes from PL #1-3 via conventional HDR, with linearized HDR templates transfected at equivalent molar amounts or 1-2 µg, respectively. (n = 4-11 healthy donors; Two-way ANOVA followed by Tukeýs multiple comparisons test). (**E**) Fold change in integration efficiency of ‘one-pot’ PASTA compared to conventional HDR with indicated mean. (n = 8 healthy donors). Asterisks represent different p-values calculated in the respective statistical tests (ns: p ≥ 0.05; *: p < 0.05; **: p < 0.01; ***: p < 0.001; ****: p < 0.0001).

### Despite similar efficiency in landing pad insertion, HDR enables higher downstream integration rates than prime editing

We next compared ‘one-pot’ PASTA to PASSIGE, which uses prime editing with two pegRNAs (Twin-PE)^23^ instead of HDR to install the landing pad. While ‘one-pot’ PASTA targets *TRAC* exon 1, PASSIGE edits the upstream intron, as described in the original publication^37^. To directly compare HDR and PE for landing pad insertion, we first assessed integration efficiency in the absence of integrase or plasmid DNA using Sanger-Sequencing coupled with TIDER (Tracking of Insertions, DEletions and Recombination events) analysis^38^. HDR-mediated insertion was dose-dependent, ranging from 25% to 95%, whereas prime editing consistently yielded ∼75% across tested pegRNA doses (2 µg vs. 6 µg) and editor variants (PEmax vs. PE7 [Ref ^28^]) (**Fig. 3A, B**). Notably, in our hands, prime editing failed when using *in vitro* transcribed pegRNAs and required synthesized and chemically protected pegRNAs (**Suppl. Fig. 3**). To compare full editing outcomes, we integrated a tricistronic payload (PL #3, as in **Fig. 2A**) delivered as nanoplasmid (NP) with minimimal bacterial backbone, using tEGFR expression as a reporter. Since tEGFR is driven by an internal EF1α promoter, its expression is independent of the exact integration site, enabling unbiased comparison between editing strategies. ‘One-pot’ PASTA achieved ∼37% tEGFR⁺ cells, while PASSIGE yielded only 5-6% (**Fig. 3C**), despite comparable landing pad insertion efficiencies (**Fig. 3A, B**). No tEGFR expression was seen in nanoplasmid-only controls, as measurements were performed ∼10 days post electroporation, by which time transient expression had resolved. These findings were consistent across donors (**Fig. 3C, D**) and were further validated by in-out droplet digital (dd)PCR analysis of extracted genomic DNA (**Fig. 3E**).

**Fig. 3.**
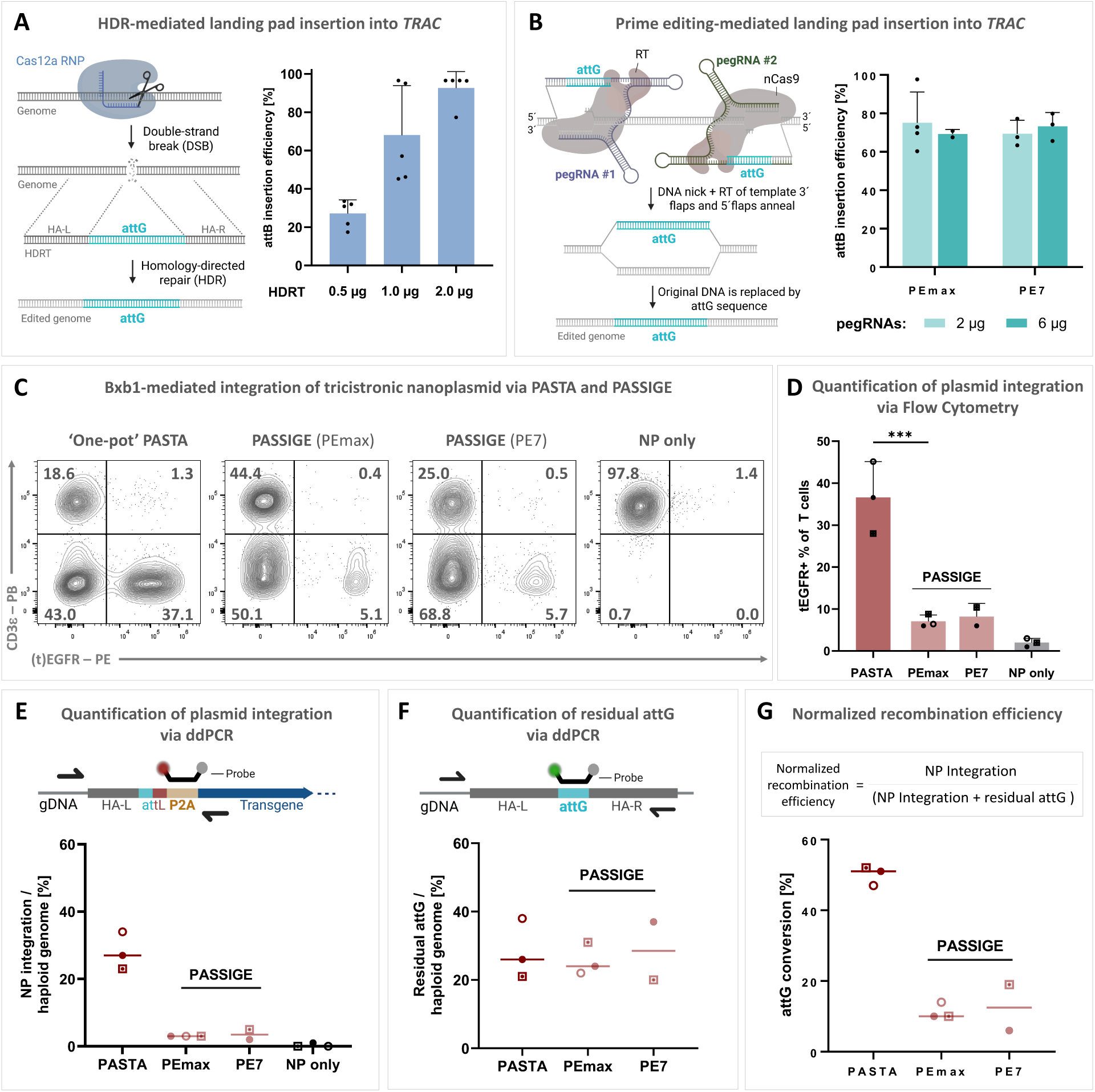
‘One-pot’ PASTA outperforms PASSIGE for large transgene insertions. (**A**) HDR-mediated insertion of an attG landing pad into exon 1 of the *TRAC* locus in primary human T cells. (**B**) Intronic attG landing pad installation via prime editing with two prime editor/pegRNA pairs (twin-PE) using either the PEmax or PE7 version of the prime editor and 2 µg or 6 µg of each pegRNA respectively. Landing pad insertion efficiencies for the HDR- and PE-mediated strategies were determined by locus-specific Sanger sequencing and TIDER analysis on genomic DNA (gDNA) isolated on day 4 post electroporation. (n = 2-3, healthy donors). (**C**) Representative flow cytometry plots for Bxb1-mediated integration of PL #3 delivered as a 7.3 kb nanoplasmid (NP, with minimalized bacterial backbone) with landing pad insertion via HDR (PASTA) or prime editing (PASSIGE) at the *TRAC* locus. Integration efficiencies were determined by flow cytometry analysis of tEGFR expression driven by the EF1a-promoter, allowing comparability between different targeted regions of *TRAC*. Flow cytometry was performed on day 10 post electroporation. (**D**) Quantification of nanoplasmid integration via PASTA and PASSIGE based on flow cytometry analysis of tEGFR expression on day 10 post electroporation. Nanoplasmid transfection alone (TF NP) served as control for transient/cytosolic tEGFR expression from the plasmid. (n = 2-3 healthy donors; One-way ANOVA followed by Šidák’s multiple comparisons test). (**E**) Quantification of nanoplasmid (NP) integration via PASTA and PASSIGE on DNA level via ddPCR performed on isolated genomic DNA. (n = 2-4 healthy donors). (**F**) ddPCR quantification of residual (inserted, but non-recombined) attG landing pads in the genome. (n = 3 healthy donors). (**G**) Normalized recombination efficiencies of Bxb1 integrase in the PASTA and PASSIGE systems, calculated from the ddPCR quantifications of nanoplasmid integration and resiudal attG in (E) and (F). (n = 2-4 healthy donors).

To determine whether the reduced efficiency observed with PASSIGE was due to inefficient landing pad insertion or impaired integrase-mediated recombination, we quantified residual attG landing pads – representing inserted but unrecombined sites – using ddPCR, which revealed similar frequencies of residual attG in both ‘one-pot’ PASTA and PASSIGE samples (**Fig. 3F**). By comparing nanoplasmid integration levels to residual attG quantities, we calculated a normalized recombination efficiency, revealing that recombination efficiency of Bxb1 within the ‘one-pot’ PASTA system was ∼5-fold higher than in PASSIGE (**Fig. 3G**). These data suggest that the recombination step, rather than landing pad insertion, is the primary limiting factor in PASSIGE.

### Engineered CAR T cells combine redundant tumor recognition with a clinically compatible safety mechanism

To evaluate individual functionality of all three payload components delivered via the tricistronic PL #3, we first tested tEGFR as a surface marker for enrichment. Magnetic bead-based selection enriched the tEGFR⁺ population from ∼37% to ∼96% (**Fig. 4A**), demonstrating its utility for downstream processing and potential use as an *in vivo* safety switch^39^. To assess CAR function, we conducted co-culture assays with wild-type Raji cells – naturally expressing both CD19 and CD20 – and engineered knockout cells lacking either one (CD19-KO, CD20-KO) or both antigens (CD19/20-DKO) (**Fig. 4B**). Activation and cytotoxicity were observed whenever at least one CAR-antigen interaction was possible, confirming that both CARs are independently functional. T cells edited with the monocistronic CD19-CAR (PL #1) served as controls and only responded to CD19-expressing targets but failed to eliminate CD19-KO tumor cells (**Fig. 4C, D**). These results confirm the activity of all three transgenes and highlight the potential of dual-antigen targeting to prevent tumor recurrence due to single-antigen loss^4^.

**Fig. 4.**
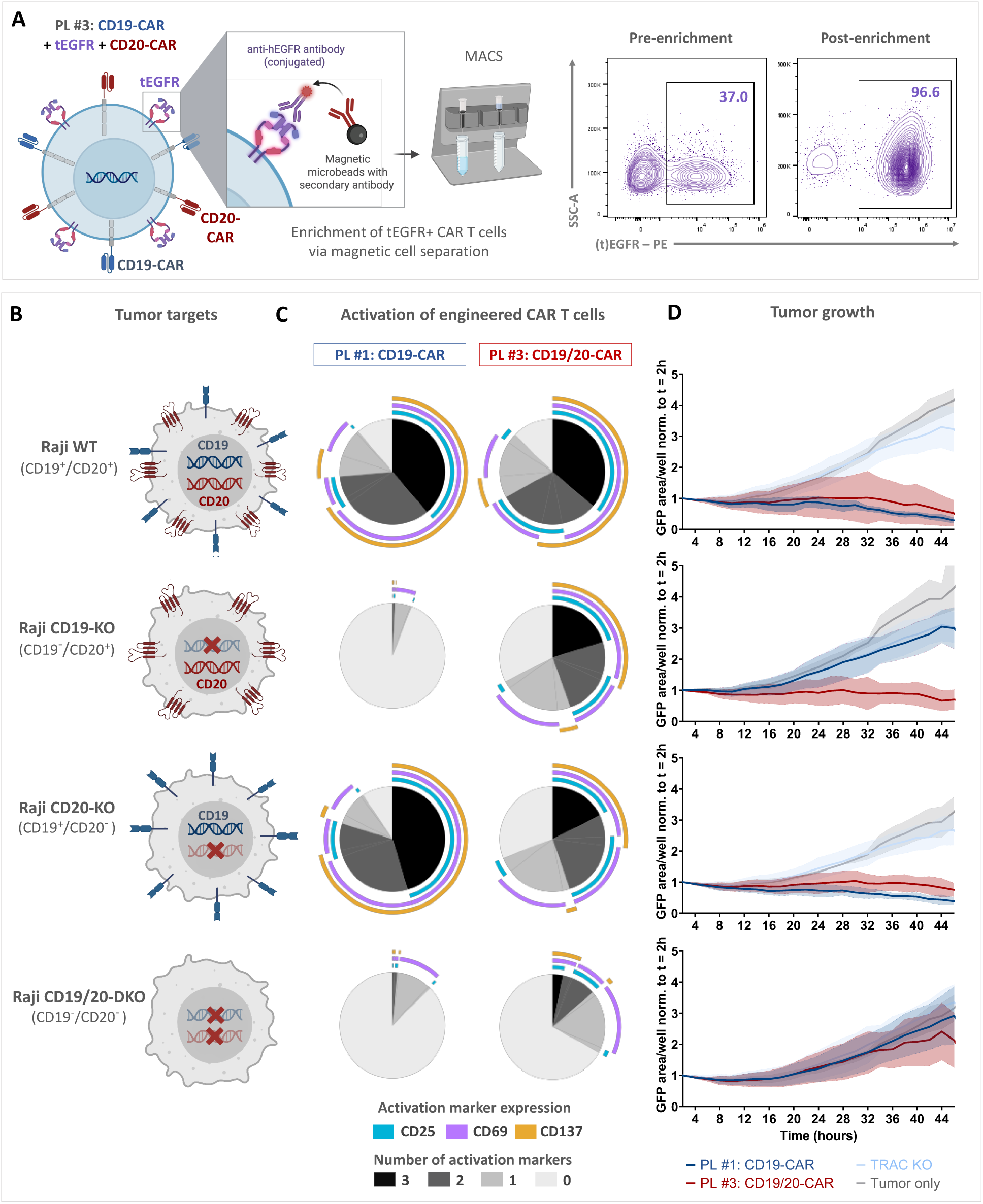
Evaluation of multi-functional CAR T cells generated with one-pot PASTA. (**A**) tEGFR enables enrichment via magnetic cell separation. A PE-conjugated antibody against human EGFR was used, followed by labeling with magnetic microbeads conjugated to secondary anti-PE antibodies. (**B**) Tumor targets for functional evaluation of CD19- and CD20-CARs. Single and double knockouts (KO / DKO) were performed on CD19/20-positive Raji (B cell lymphoma) cells to assess cytotoxic activity of CAR T cells with mono- or bispecific targeting of CD19 and CD20. (**C**) Activation of CAR T cells after 24h of co-culure with the Raji cell lines. CAR T cells with dual-specificity for CD19 and CD20 (generated by integration of PL #3) were compared against CD19-directed monospecific CAR T cells (generated by integration of PL #1). (**D**) Tumor growth of the Raji cell lines in a 1:1 co-culture with the of mono- or bispecific CAR T cells monitored over 48 hours using the Incucyte live-cell analysis system. In (C) and (D), Raji cells were co-cultured with mono- or bispecific CAR T cells at an effector-to-target (E:T) ratio of 1:1. (n = 4-5 healthy donors).

### Engineered integrases and nanoplasmids enhance the efficiency of ‘one-pot’ PASTA

Our findings suggest that the overall efficacy ‘one-pot’ PASTA is primarily limited by the efficiency of serine integrase-mediated recombination, which depends on the enzyme and the availability of its substrate. We hypothesized that integration rates might be further improved by using alternative integrases to Bxb1, the current gold-standard in the field. To evaluate this in T cells, we developed a reporter system for parallel screening of four serine integrases – Bxb1, PhiC31 and the recently discovered LSRs Pa01 and Kp03^40^. An attG landing pad array (with attachment sites for all four integrases) was inserted into the *GAPDH* locus^41^. As the landing pad array inevitably contains stop codons, we incorporated an artificial intron^42^ followed by a splice acceptor and a GFP reporter in the HDR template (**Fig. 5A, Suppl. Fig. 4A**). A corresponding recombination plasmid carried the matching attV array upstream of a splice acceptor and an RFP reporter cassette. Successful HDR resulted in GFP⁺ cells, while LSR-mediated recombination led to RFP expression (**Fig. 5B, Suppl. Fig. 4B**).

**Fig. 5.**
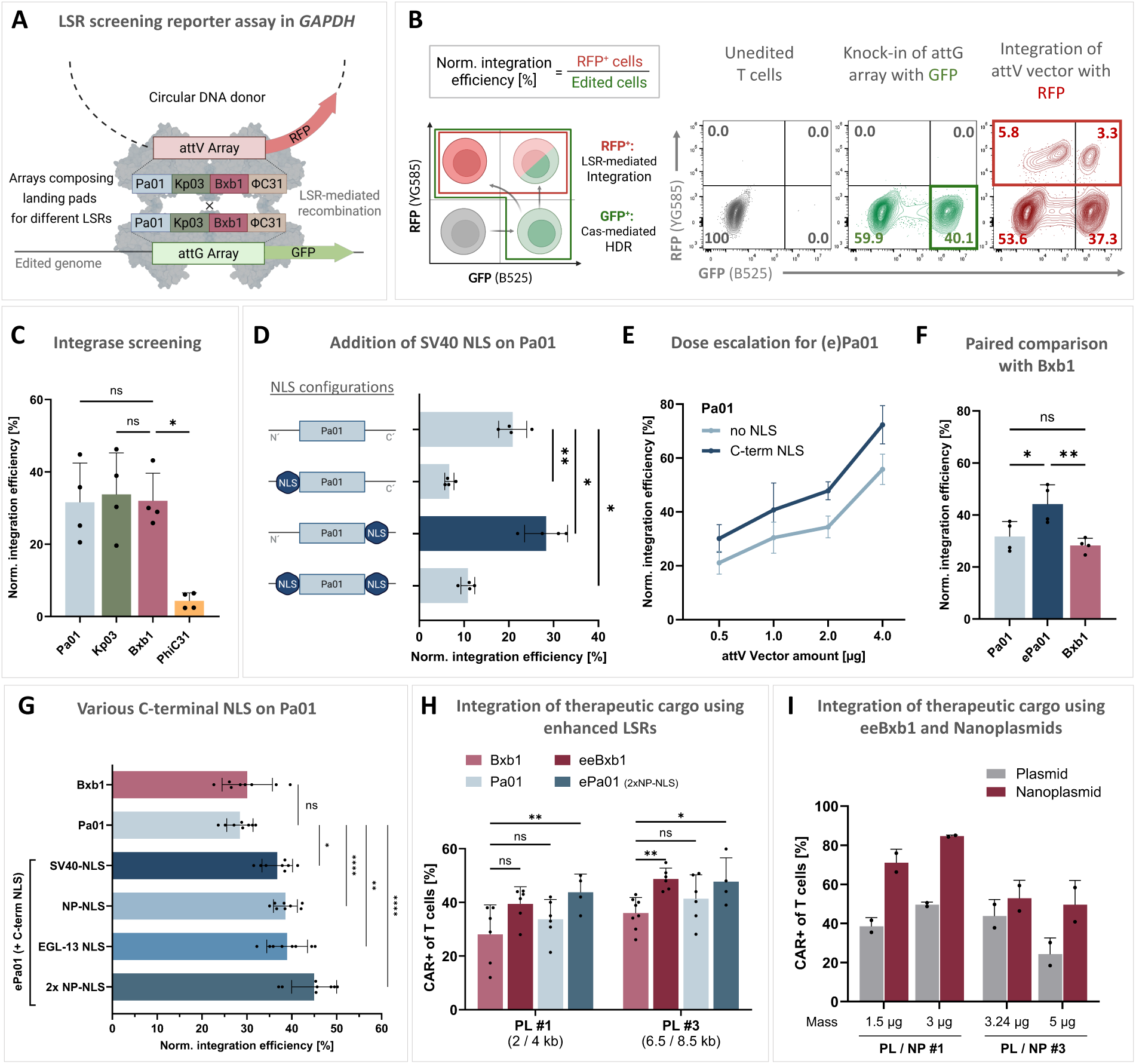
Enhancing ‘one-pot’ PASTA with engineered integrase ePa01. (**A**) Schematic of integrase screening assay in the *GAPDH* locus of primary human T cells. The reporter-based screening assays comprises an array of four attG landing pads for the large serine integrases Pa01, Kp03, Bxb1 and PhiC31 tagged with GFP for HDR-mediated insertion into the *GAPDH* locus and a circular plasmid containg the corresponding attV landing pads and RFP to be expressed upon successful recombination into the genome. (**B**) Quantification of LSR-mediated recombination efficiency by normalizing RFP-positive cells to total edited cells (sum of GFP-positive and RFP-positive cells) (left). Representative flow cytometry plots for GFP and RFP expression, showing different editing outcomes using the reporter assay (right). (**C**) Normalized integration efficiences of the integrases Pa01, Kp03, Bxb1 and PhiC31. (n = 4 healthy donors; One-way ANOVA followed by Tukeýs multiple comparisons test). (**D**) Evaluation of an SV40 nuclear localization signal (SV40-NLS) fused to Pa01 in various configurations – C-terminal, N-terminal and on both termin – for its effect on integrase activity. (n = 4 healthy donors; One-way ANOVA followed by Tukeýs multiple comparisons test). (**E**) Normalized integration efficiencies of wild-type Pa01 and Pa01 with C-terminal SV40-NLS across increasing plasmid doses. (n = 4 healthy donors). (**F**) Normalized integration efficiencies of wild-type Pa01, enhanced Pa01 (ePa01) with C-terminal SV40-NLS and Bxb1 in paired experiments at a plasmid dose of 1 µg. (n= 4 healthy donors; One-way ANOVA followed by Tukeýs multiple comparisons test). (**G**) Comparison of different C-terminally fused NLS regarding their effect on Pa01 activity. (n = 4 healthy donors; One-way ANOVA followed by Tukeýs multiple comparisons test). (**H**) Integration of PL #1 and PL #3 as therapeutic cargo into the *TRAC* locus using Bxb1, the previously described phage-evolved eeBxb1, wild-type Pa01 and ePa01 with two C-terminally fused NP-NLS motifs. PL #1 and #3 were delivered in equimolar amounts. Integration efficiencies were assessed by flow cyotmetry analysis of CAR expression on day 4 after electroporation. (n = 4-8 healthy donors; Two-way ANOVA with Šidák’s multiple comparisons test). (**I**) Efficiencies of eeBxb1-mediated integration of PL #1 and #3 delivered either as standard plasmids (PL) or as nanoplasmids (NP) with minimalized bacterial backbone. (n = 2 healthy donors).

Among the integrases tested, PhiC31 exhibited minimal activity, while Pa01 and Kp03 performed on par with the standard Bxb1 integrase (**Fig. 5C**). Notably, PhiC31 and Bxb1 have previously been engineered to harbor nuclear localization signals (NLS) for improved activity in human cells^23,43–45^. In contrast, Pa01 and Kp03 integrases lacked NLS, offering a potential avenue for further optimization. We focused on Pa01 due to its higher specificity within the human genome^40^, and systematically evaluated different NLS configurations. Addition of a single C-terminal SV40 NLS significantly boosted Pa01’s activity, whereas N-terminal NLS placements impaired its function (**Fig. 5D**). The NLS-enhanced variant exhibited increased recombination efficiency over wild-type Pa01 across different substrate dose levels (**Fig. 5E**) and also significantly outperformed Bxb1. (**Fig. 5F**). Further screening of different NLS resulted in the selection of two nucleoplasmin (NP) NLS at the C-terminus, yielding our best-performing enhanced (e)Pa01 variant to date (**Fig. 5G**).

To compare the performance of wildtype and engineered integrases in a therapeutically relevant setting, we targeted monocistronic (PL #1) and tricistronic (PL #3) payloads to the *TRAC* locus. We compared our optimized ePa01 and the recently described evolved and engineered Bxb1 (eeBxb1)^37^ against their wildtype versions. Both engineered integrases outperformed their corresponding wild-type counterparts and achieved comparable integration efficiencies when used in the ‘one-pot’ PASTA approach (**Fig. 5H**). Finally, we used this therapeutically relevant setup to compare donor templates by delivering conventional plasmids versus nanoplasmids – minimimal-backbone DNA vectors. Nanoplasmids consistently improved integration efficiency with eeBxb1 for both payload sizes, increasing monocistronic integration from 50% to 80%, and integration of the larger tricistronic payload from 25% to 45% (**Fig. 5I**). Additionally, we observed that nanoplasmids improved transgene expression and viability in engineered T cells (**Suppl. Fig. 5**). Consequently, engineered LSRs and plasmids with minimal bacterial backbones provide promising tools to advance ‘one-pot’ PASTA for clinically scaled manufacturing of engineered CAR T cells.

## Discussion

Our study introduces ‘one-pot’ PASTA, a non-viral platform for programmable and site-specific integration of large, multi-cistronic transgenes into primary human T cells. By combining HDR to install a short landing pad and subsequent LSR-mediated recombination, ‘one-pot’ PASTA enables efficient, targeted insertion of genetic payloads exceeding the size limits of conventional HDR. We demonstrate the platform’s versatility across multiple genomic loci, its capacity to stably integrate complex dual CAR constructs, and superior efficiency over prime-editing–based strategies, such as PASSIGE, in primary human T cells. Finally, we show that ‘one-pot’ PASTA’ can be boosted using engineered and evolved serine integrases, such as ePa01 or eeBxb1, and circular DNA with minimal backbones. To support broad accessibility, a set of core PASTA plasmids have been deposited with **Addgene**.

In addition to outperforming conventional HDR for large gene transfer, ‘one-pot’ PASTA offers distinct advantages over viral vectors and undirected transposases. While randomly integrating lenti- or retroviral vectors are the current gold standard in clinical CAR T cell engineering, their efficiency declines markedly with transgenes exceeding ∼7kb due to reduced viral titers and impaired genomic integration^46,47^. In contrast, ‘one-pot’ PASTA offers precision through site-specific integration and greater scalability, as it relies on circular non-viral DNA vectors, which are not subject to similar size constraints. This precision reduces the risk of insertional mutagenesis associated with retroviruses and transposases^10,11^, and expands the possibilities for T cell engineering. For instance endogenous loci can be exploited to modulate transgene expression levels (**Fig. 1**)^14^, enabling selection of edited cells (e.g., via “SLEEK” at *GAPDH* locus^41^), and simultaneously knocking out genes such as the TCR/CD3 complex to reduce risks like graft-versus-host disease in allogeneic applications^16,48,49^. As a proof of concept, we demonstrated ‘one-pot’ PASTA’s capability to integrate complex payloads into the *TRAC* locus, including two CARs, an additional promoter, and a tEGFR module, which serves as a marker for cell enrichment, tracking, and potentially as a safety switch^39^. Functionally, such a dual-CAR T cell product proved effective in co-cultures with CD19-/ CD20-KO Raji cells, highlighting its potential in heterogeneous malignancies like acute B cell leukemia and aggressive B cell lymphoma, where antigen-negative relapse remains a significant challenge^4^. Moving forward, substantial efforts should be directed toward identifying the optimal CAR constructs and additional transgenes to enhance therapeutic potency and broaden clinical applicability. Herein, combining pooled transgene screens with ‘one-pot’ PASTA ^50,51^ could accelerate the development of next-generation CAR constructs and even programmable synthetic circuits (e.g. incorporation of inducible systems).

Our data demonstrate that ‘one-pot’ PASTA achieves higher recombination efficiency in T cells compared to PASSIGE (**Fig. 3G**), despite employing an optimized Twin-PE strategy^37^ and chemically protected pegRNAs^28,29^. HDR and Twin-PE may differ in their kinetics for landing pad insertion. Delayed completion of the editing process with Twin-PE could explain the lower recombination efficiency of PASSIGE, either through blocking of the landing pad site by a PE-DNA-complex or dilution of the integrase and its substrate in proliferating cells. The difference may also be related to the LSR-mediated recombination step: during PASSIGE, the pegRNA is unlikely to serve as a substrate for the integrase, requiring the insertion of the landing pad into the genome first, to enable subsequent integration of the plasmid DNA. In contrast, ‘one-pot’ PASTA may allow LSR-mediated recombination of the plasmid and the HDRT prior to insertion into the genome. An unresolved integrase-HDRT-plasmid complex could adopt a tertiary structure that facilitates integration more efficiently than a conventional HDR donor. Elucidating the exact underlying mechanisms could inspire further improvements of ‘one-pot’ PASTA.

The simplicity of ‘one-pot’ PASTA compared to strategies like PASSIGE and PASTE likely enables broader accessibility of LSR-mediated gene transfer in HDR-competent cells. Laboratories already familiar with HDR can implement the system, as it relies on commonly available tools like conventional Cas nucleases, guide RNAs and short synthetic HDR templates. A related method, ‘ONE-STEP tagging,’ was recently demonstrated in immortalized and pluripotent cell lines and, to a limited extent, in primary T cells, using plasmid-based nuclease/integrase expression and single-stranded oligodeoxynucleotides (ssODN) landing pad donor. While the emergence of both strategies underscores the broad relevance of combining HDR with integrases, PASTA implements this principle through delivery of Cas-RNPs and integrase mRNA, achieving efficient integration of large payloads in primary T cells^52^. While PE-alternatives, such as PASSIGE, may offer advantages when reprogramming non-dividing cells, HDR-guided integrase recombination represents a more straightforward, efficient and potentially more versatile approach for T cells and other HDR-competent cell types that are amenable to plasmid transfection.

The performance of the LSR and the availability of circular DNA substrates constrain the efficiency of ‘one-pot’ PASTA and related systems such as PASSIGE. Addition of C-terminal NLS to Pa01 significantly enhanced its recombination efficiency, on par with eeBxb1, which has undergone extensive directed evolution^37,53^. The newly developed ePa01 is an attractive candidate to be paired with PE or other templated insertion tools, such as CLICK-editors^54^, to reprogram non-dividing cells or cells that are not amenable to HDR.

Beyond enzyme optimization, maximizing the quality and delivery of circular DNA templates is crucial. The use of nanoplasmids or other minimal-backbone plasmids can improve delivery efficiency while reducing cellular toxicity (**Fig. 5, Suppl. Fig. 5**) ^12,13^. Our results suggest that a co-integrated plasmid backbone can affect transgene expression from endogenous loci (**Suppl. Fig. 5A, B**). To avoid integration of bacterial sequences, ‘one-pot’ PASTA could also be adapted to facilitate recombination mediated cassette exchange^55^, thereby allowing precise replacement of genomic elements without backbone carryover.

Overall, ‘one-pot’ PASTA represents a versatile, scalable, and site-specific strategy for engineering primary human T cells, with significant potential in both therapeutic development and basic research.

## Methods

### Ethics statements

The study with material from human participants was performed in accordance with the declaration of Helsinki (Charité ethics committee approval EA4/091/19 and EA1/052/22, Baylor College of Medicine IRB approval H-45017).

### Cell culture

Peripheral blood mononuclear cells (PBMCs) were isolated by density gradient centrifugation using Ficoll and subsequently enriched for CD3⁺ T cells via magnetic-activated cell sorting (MACS), as previously described^16,18^. T cells were cultured in antibiotic-free CTL medium (1:1 mixture of Advanced RPMI and Click’s medium supplemented with 10% FCS and 1% Glutamax) in the presence of IL-7 (10 ng/ml, Miltenyi Biotec) and IL-15 (5 ng/ml, Miltenyi Biotec). After isolation, T cells were activated on αCD3/CD28-coated tissue culture plates for 2 days.

### Cloning

Plasmids were assembled via In-Fusion cloning (Takara) at half of the manufacturer’s recommended volumes to reduce costs per reaction. All plasmids were sequence-verified via whole plasmid next generation sequencing (Eurofins Genomics or Plasmidosaurus).

### HDRT generation

HDRTs were prepared as previously described^14,16^. Briefly, the DNA templates were amplified from plasmids by PCR using the KAPA HotStart 2x Readymix (Roche). The PCR products were then purified using the AMPure XP paramagnetic beads (Beckman Coulter Genomics). For this, the DNA was mixed in a 1:1 ratio with the beads, incubated at room temperature for 10 min, and then placed on a DynaMag-2 stand (Invitrogen, Thermo Fisher Scientific) for 10 min. Next, the DNA-bead mix was washed two times under sterile conditions with 70% ethanol. Finally, the DNA was eluted in 3 µl of nuclease-free water per 100 µl PCR product. Concentrations were determined using the Nanodrop ND-1000 spectrophotometer (Thermo Fisher Scientific) and adjusted to 2 µg/µl.

### Preparation of circular plasmid DNA for integrase-mediated recombination

Plasmids containing the transgene payload of interest were transformed into stellar competent *E. coli* and cultured overnight at 37°C in 200-500 ml of LB broth supplemented with ampicillin. The plasmids were isolated using the ZymoPURE II Plasmid Maxiprep Kit (Zymo Research) according to the manufactureŕs protocol. Subsequently, isolated plasmid DNA was concentrated using AMPure XP beads as previously described to reach desired concentrations of 2-7 µg/µl. Nanoplasmids were produced from original plasmid sequences by Aldevron.

### mRNA generation

DNA sequences encoding the integrases were separately cloned into a plasmid backbone containing a mutated dead (d)T7 promoter, 5′ UTR, Kozak sequence, and 3′ UTR. Prime editor constructs were sourced from Addgene: pT7-PEmax for IVT was a gift from David Liu (Addgene plasmid # 178113) and pT7-PE7 for IVT was a gift from Brittany Adamson (Addgene plasmid # 214813). eeBxb1 was a gift from David Liu (Addgene plasmid # 222339). The Pa01 integrases were sourced from Durrant et al^56^. The dT7 promoter was corrected during PCR amplification, which also introduced a 120 bp poly(A) tail to enhance mRNA stability. *In vitro* transcription was performed as previously described^31,57^ using the HiScribe™ T7 High Yield RNA Synthesis Kit (NEB) with N1-methyl-pseudouridine (Trilink) in place of uridine and co-transcriptional CleanCap AG (Trilink). *In vitro* transcription of pegRNA was performed without CleanCap and modified bases as previously described^58^. DNase I treatment was used to degrade template DNA, and the resulting RNA was purified using the Monarch^®^ RNA Cleanup Kit (NEB). mRNA quality was assessed by denaturing agarose gel electrophoresis and quantified via the Qubit 4 Fluorometer (Thermo Fisher Scientific) using the ssRNA BR assay. Transcripts were stored at –80 °C, and 1µg was used per electroporation.

### Genome editing and Electroporation

Two days after cell isolation, electroporation was performed using the Lonza 4D Nucleofector system (16-well strips, 20 µl volume). The volumes and conditions described refer to the smallest possible electroporation format (20 µl total volume) and are compatible with linear upscaling in larger electroporation cuvettes. Cas-RNPs were preassembled by incubating 0.5 µl poly (L-glutamic acid) (100 µg/µl, mol wt 15,000-50,000, Sigma-Aldrich), 0.48 µl of chemically modified guide RNA (100 µM) (IDT), and 0.4 µl of recombinant Cas protein (Alt-R SpCas9 V3, conc. = 10 µg/µl) (IDT) or Cas12a (Alt-R Cas12a Ultra, conc. = 10 µg/µl) (IDT) for 15 min at room temperature, as described previously^15,33^. 1.0-1.5×10^6^ activated T cells were harvested, washed twice with PBS and resuspended in P3 buffer. Meanwhile, the reagents for one-pot PASTA were premixed in 96-well PCR plates in the following order: 1.38 µl Cas RNP, 0.5 µl linear HDR template (2 µg/µl), up to 2 µl circular plasmid DNA (>2 µg/µl), and 0.5 µl integrase mRNA (2 µg/µl). Finally, 20 µl of cells in P3 buffer were added, gently mixed, and transferred into 16-well electroporation strips using multichannel pipettes. To eliminate air bubbles before electroporation, the strips were tapped firmly on the benchtop. Electroporation was performed using programs EH-115 or EO-115. Immediately after electroporation, 100 µl of pre-warmed T cell culture medium was added per well and incubated for 10 min at 37°C. Cells were then transferred to 96-well U-bottom plates containing media supplemented with HDR-enhancing compounds – a combination of 0.25 µM AZD7648 and 1 µM ART558 (MedChemExpress)^35,36^. Medium was changed the next day to minimize prolonged exposure to DNA repair inhibitors.

### Prime editing

For experiments involving twin-prime editing for landing pad installation into the genome, electroporation using the Lonza 4D Nucleofector system was performed, as described above. In these experiments, the Cas-RNP and HDR template were replaced with 2 µg of in vitro-transcribed prime editor mRNA (either PEmax or PE7) and 2–6 µg of each synthesized and chemically modified pegRNA (IDT), which encoded the landing pad sequence for reverse transcription. For PASSIGE, 2 µg of each pegRNA was used and co-transfected with integrase mRNA and circular plasmid DNA in the amounts as described above.

### Flow cytometry

Transgene expression, cytotoxicity, and T cell activation were assessed using a Cytoflex LX flow cytometer (Beckman Coulter) in 96-well plate format. Antibody panels are detailed in **Supplementary Table 2** and staining procedures were performed as previously described^16^. Representative gating strategies are provided in **Supplementary Figure 6**.

### Quantification of genomic landing pad insertion

For quantification of genomic landing pad insertion into *TRAC* via HDR and prime editing, genomic DNA was isolated on day 4 post electroporation using the DNeasy Blood & Tissue Kit (Qiagen). A locus-specific PCR for amplification of the target region was performed and purified PCR fragments (DNA Clean & Concentrator-5 kit, Zymo Research) were sent for Sanger sequencing (LGC Genomics). The editing efficiency was determined using TIDER^38^ to quantify attG landing pad insertion.

### Generation of Raji cell lines

Raji cell lines were electroporated with Cas9-gRNA complexes targeting CD19 and CD20 using program DS-104, as previously described^59^. Single-cell clones were generated by limiting dilution in 96-well plates and screened to identify clones with complete (100%) knockout. Subsequently, cells were lentivirally transduced to express nuclear GFP and subsequently sorted for GFP positivity.

### T cell activation assay

Activation of generated CAR T cells was measured upon stimulation with tumor cells. CAR T cells were expanded for 14 days following electroporation and subsequently rested for 24 hours in cytokine-free medium prior to the assay. A total of 1 × 10^5^ T cells were seeded into v-bottom 96-well plates and co-cultured in a 1:1 target-to-effector cell ratio with GFP-labeled Raji target cells. After 24 hours of co-culture, cells were stained for CD25, CD69 and CD137 as activation markers (**Suppl. Table 2**).

### Co-culture assays via live cell imaging

In vitro tumor control of generated CAR T cells was assessed via live cell imaging of GFP-expressing cancer cells on an Incucyte^®^ device (Sartorius). CAR T cells were expanded for 14 days following electroporation and subsequently rested for 24 hours in cytokine-free medium prior to the assay. A total of 2.5 × 10^4^ CAR^+^ T cells were seeded into a flat-bottom 96-well plates and co-cultured in a 1:1 target-to-effector cell ratio with GFP-labeled Raji target cells. To account for differences in initial editing efficiencies, all effector samples were adjusted to the same frequency of CAR^+^ T cells with *TRAC*-KO cells, ensuring an equal total cell number across conditions. The plates were then placed into the Incucyte^®^ device and a repeat scanning was performed over 48 hours with scans taken every 2 hours, capturing three images per well.

### Digital droplet polymerase chain reaction (ddPCR)

ddPCR assays were designed for quantification of genomic landing pad insertion and integration of circular DNA vectors in the *TRAC* locus (primers and probes are listed in **Supplementary Table 1, ddPCR Assays**). A ddPCR assay for the RPP30 gene was used as reference^31^. Genomic DNA (gDNA) was isolated using the DNeasy Blood & Tissue Kit (Qiagen) according to the manufacturer’s instructions. 75 ng of gDNA was used as a template for a 22 µl PCR reaction containing 1.1 µl (10 µM) of each forward and reverse primer for both target and reference genes, 1.1 µl (5 µM) of each target and reference probe, 5.5 µl of 4X MultiplexMix (BioRad) and nuclease-free water. The results were analyzed with the QX Manager Software (BioRad). Genomic landing pad insertion and plasmid integration rates were quantified per haploid genome by calculating the ratio of target molecule concentration to reference molecule concentration, respectively.

### Data analysis, statistics and presentation

Flow cytometry data was analysed with FlowJo Software (BD). Prism 9 (GraphPad) was used to create graphs and perform statistics. Illustrations were created on BioRender.com.

## Supporting information

Suppl. Table 1 - Sequences

Suppl. Table 2 - Flow Cytometry Panels

## Data Availability

All new DNA and RNA construct sequences as well as primers and ddPCR probes used in this study are provided in **Supplementary Table 1**, antibody information and panel configurations in **Suppl. Table 2**. Selected plasmids created in this study will be made available via Addgene: (1) the plasmids encoding the *TRAC*-Bxb1.attG or the *TRAC*-Pa01.attG landing pad HDR-template (Addgene plasmids #247037, #247039), (2) the promoterless integration plasmids carrying the CD19-CAR with a Bxb1 or a Pa01 docking site (Addgene plasmids #247038 and #247040), and (3) the plasmid encoding for the NLS-enhanced Pa01 (ePa01) integrase used for *in vitro* mRNA transcription (Addgene plasmid #247041). These resources are shared to encourage broad adoption of this modular and easy-to-implement platform for large transgene integration in primary human T cells. All other data can be obtained from the corresponding author upon reasonable request.

## Acknowledgements

The authors would like to thank Daniel Gao and David Liu (Broad Institute) for guidance on design and sourcing of synthesized and chemically modified pegRNA for prime editing. Funding: This project was initially supported by Berlin Institute of Health Center for Regenerative Therapies, New Crossfield Grant (2023-2024) by D.L.W. and D.M.I.; I.K., J.K. and D.L.W. were supported by the SPARK-BIH program by the Berlin Institute of Health, Germany. Ibrahim laboratory is supported by an ERC Starting Grant SYNREG (101076709). This project has received funding from the European Union under Grant Agreement Nr. 101057438 (geneTIGA: genetiga-horizon.eu) to D.L.W. Views and opinions expressed are however those of the author(s) only and do not necessarily reflect those of the European Union or the European Health and Digital Executive Agency (HADEA). Neither the European Union nor the granting authority can be held responsible for them.

## Author contributions

I.K. and J.K. designed parts of the study, planned and performed experiments, analyzed results, interpreted the data, and wrote the manuscript. A.M.N. planned and performed experiments, analyzed results, interpreted the data, edited the manuscript. V.G., La.H., performed experiments, analyzed results and edited the manuscript. Y.P., Lu.H., R.K., S.S., M.S. performed experiments and analyzed results. A.R. performed experiments, provided reagents and edited the manuscript. M.P. planned experiments, analyzed results and interpreted the data. D.M.I. designed the initial parts of the study, interpreted data, provided reagents and edited the manuscript. D.L.W. designed and led the study, planned experiments, interpreted data, and wrote the manuscript. All authors reviewed, commented, and approved the manuscript in its final form.

## Supplementary Figures

**Suppl. Fig. 1.**
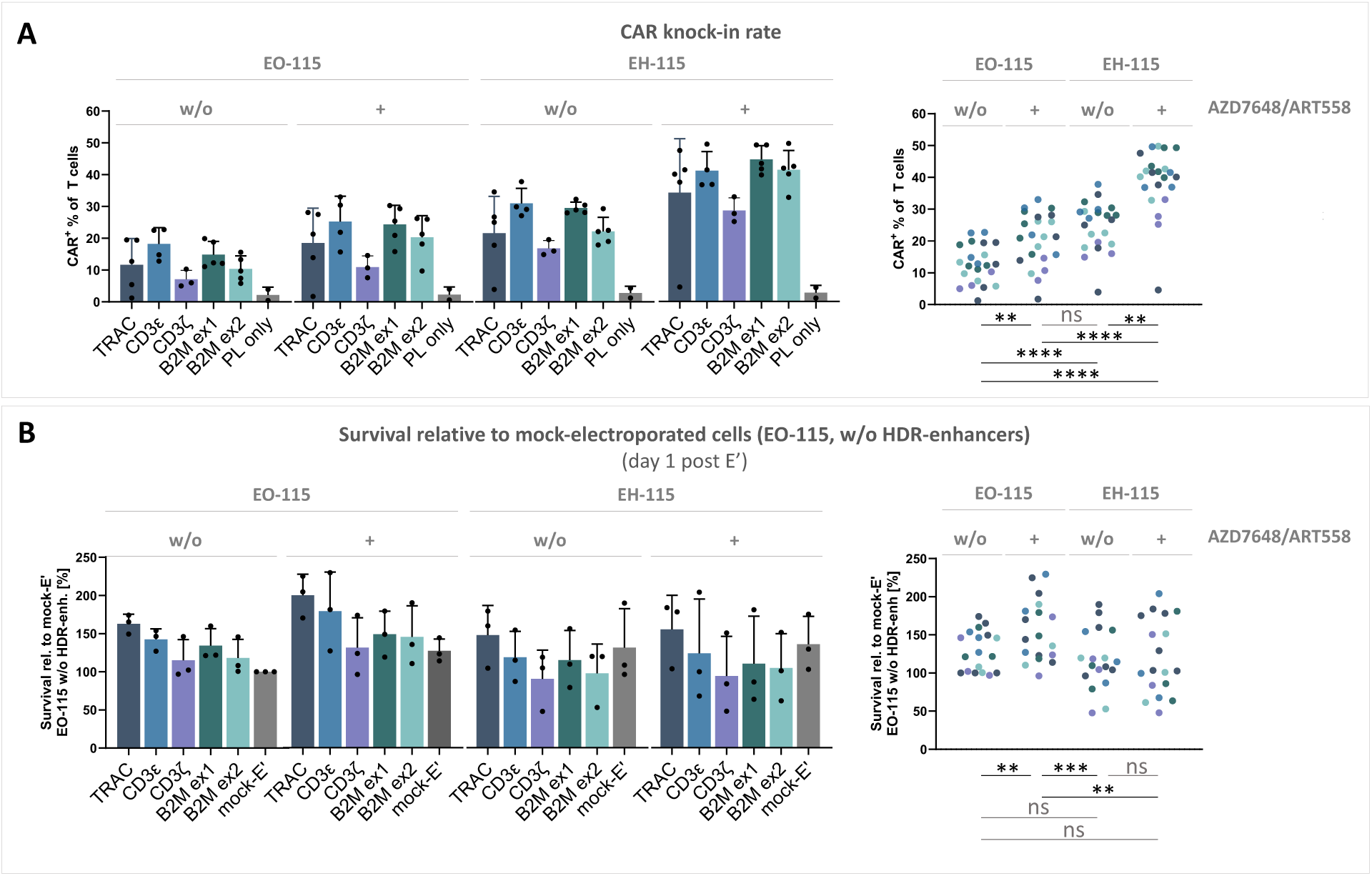
One-pot PASTA – electroporation programs and HDR-enhancers. On a Lonza 4D Nucleofector, two electroporation programs (EO-115 and EH-115) were tested side-by-side for one-pot PASTA. Following electroporation and a 10-minute resting phase in T cell media (TCM), cells were cultured either in standard TCM or in TCM supplemented with a combination of two DNA repair-modifying agents (0.25 µM AZD7648 and 1 µM ART558) (n = 2–5 healthy donors; Friedman’s test followed by Dunn’s multiple comparison test). (**A**) CD19-CAR knock-in efficiency. (**B**) Cell survival relative to mock-electroporation using EO-115.

**Suppl. Fig. 2.**
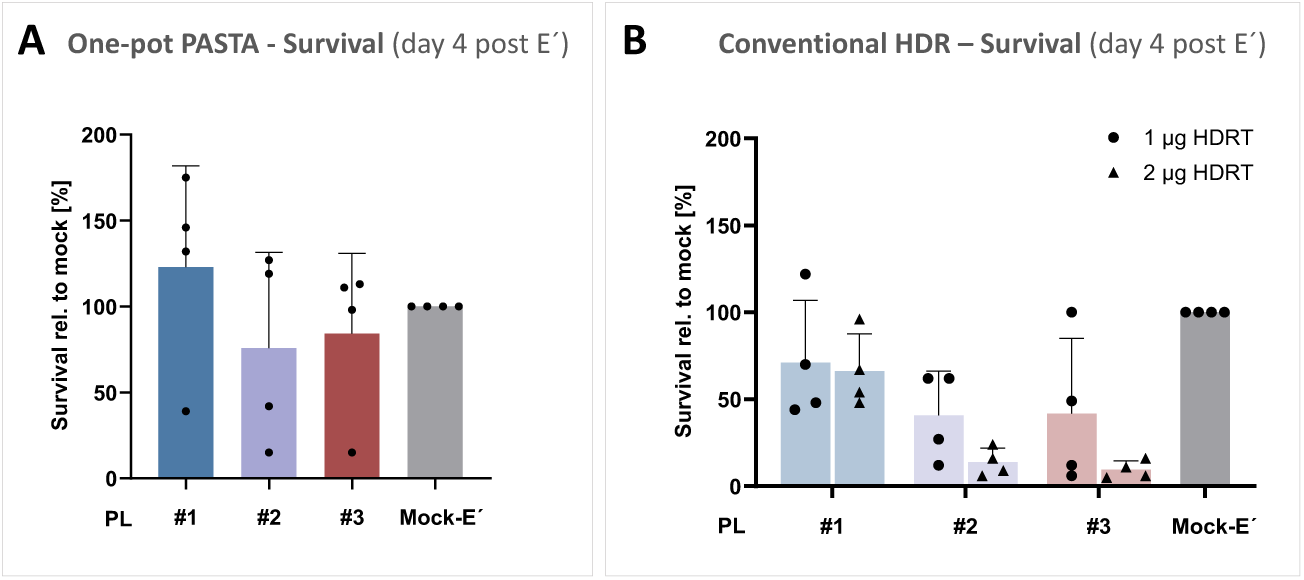
Cell survival post editing with ‘one-pot’ PASTA compared to conventional HDR. Cell survival of edited cells relative to mock-electroporation. Survival was assessed in n = 4 healthy donors for integration of mono-, bi- and tricistronic genetic payloads PL #1-3 via (A) ‘one-pot’ PASTA or (B) conventional HDR on day 4 after electroporation.

**Suppl. Fig. 3.**
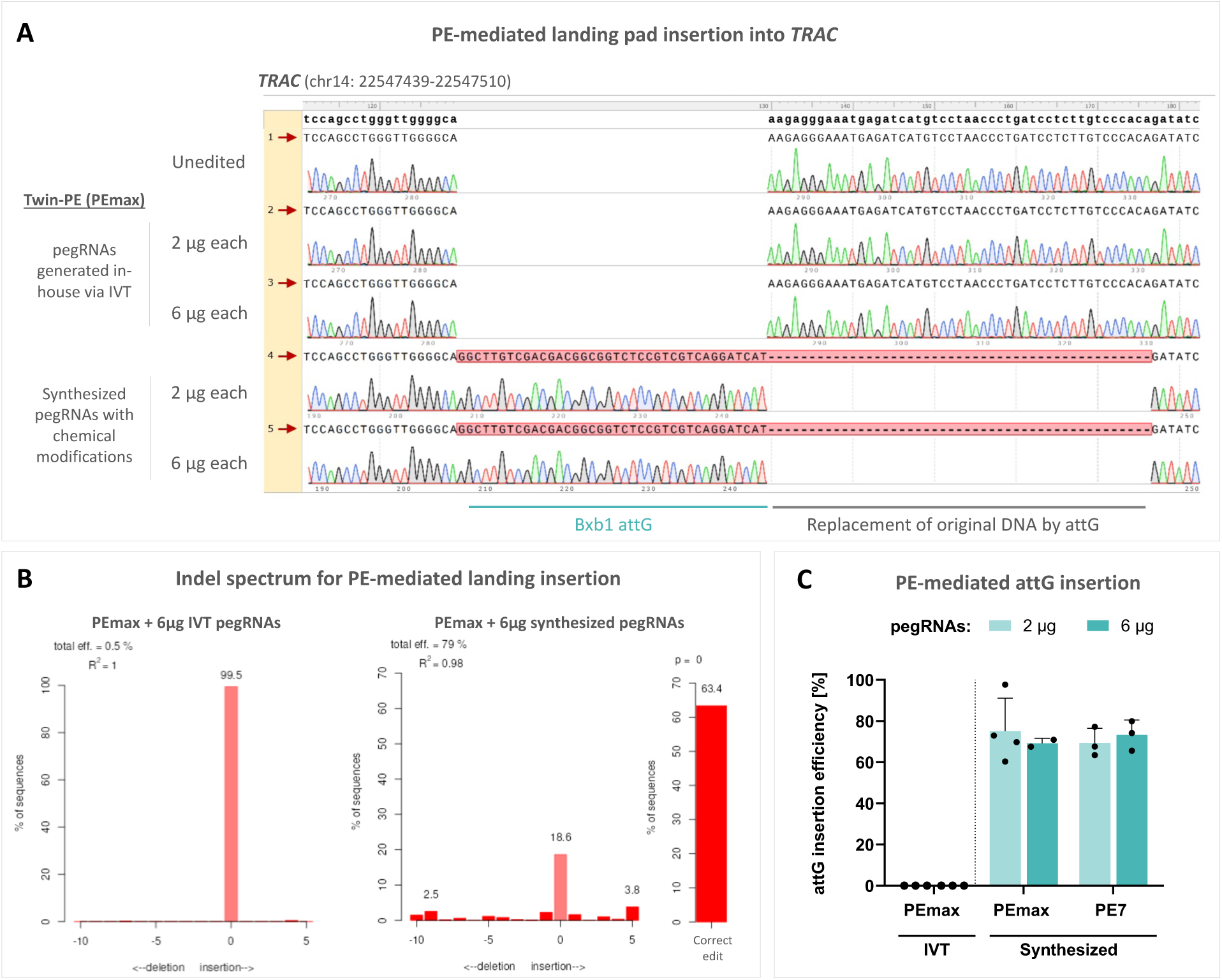
Prime editing requires chemically modified pegRNAs. (**A**) Sanger sequencing traces aligned to the *TRAC* locus at the targeted site for attG landing pad via prime editing. Chromatograms are shown for the unedited genome as reference (lane 1), prime editing using in-house generated in vitro-transcribed (IVT) pegRNAs (lanes 2 and 3) and prime editing using synthesized and chemically modified pegRNAs (lanes 4 and 5). Sequencing was performed after locus-specific amplification of genomic DNA. (**B**) Representative indel spectra for prime editing using IVT pegRNAs (left) and synthesized pegRNAs (right) determined by TIDER analysis of sequenced gDNA. (**C**) Efficiency of PE-mediated attG landing pad insertion using IVT pegRNAs versus synthesized and chemically modified pegRNAs, quantified by TIDER analysis of Sanger sequenced gDNA amplicons. (n = 2-3 healthy donors).

**Suppl. Fig. 4.**
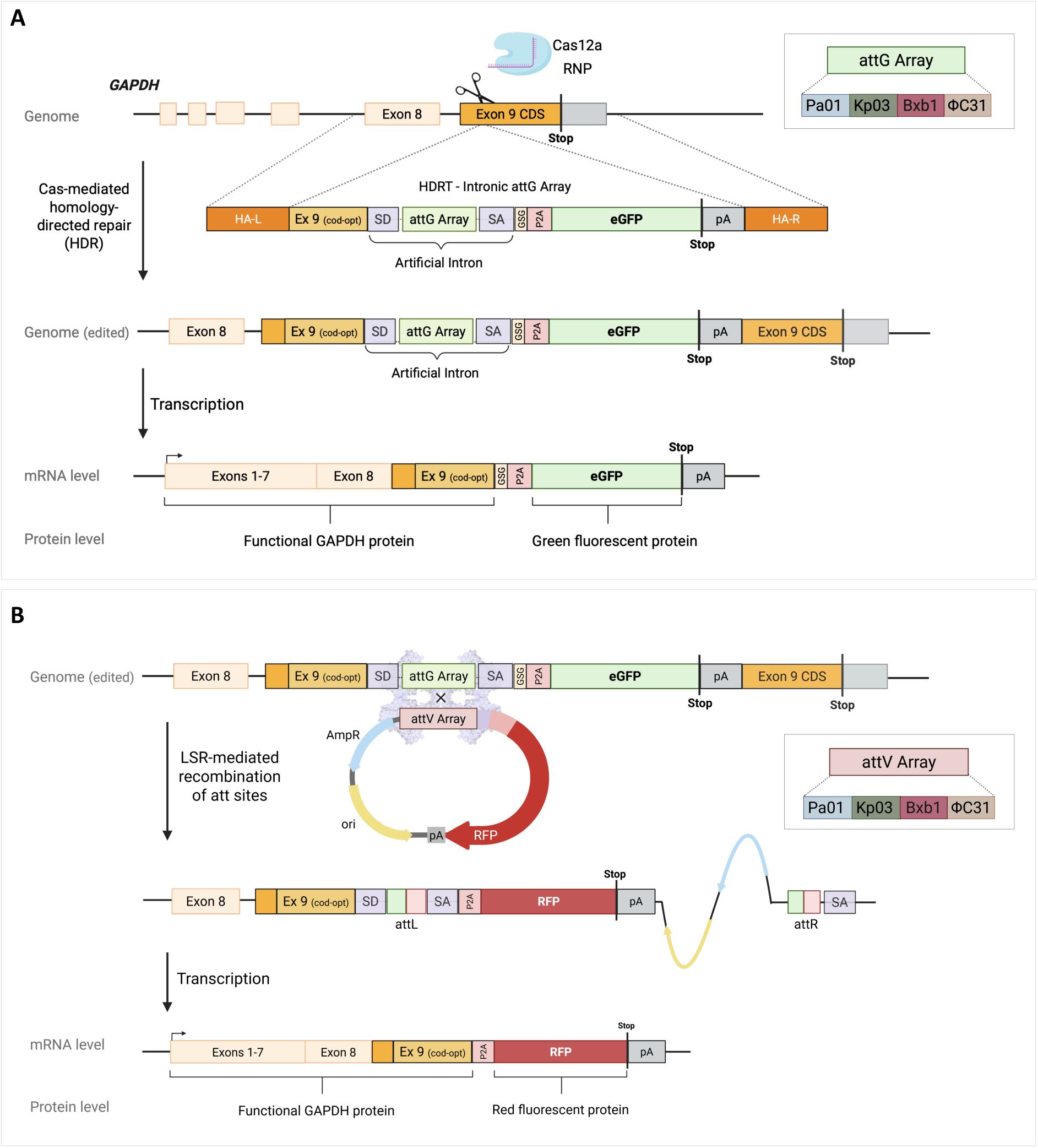
Reporter assay strategy for screening large serine integrases in the *GAPDH* locus. (**A**) Schematic showing insertion of an attG landing pad array into the last exon of *GAPDH*. The HDR template contains an array of attG landing pads for the integrases Pa01, Kp03, Bxb1 and PhiC31. The array is flanked by splice sites to create an artificial intron and is followed by a downstream eGFP reporter. Restoration of a functional GAPDH protein is achieved by in-frame integration of the recoded last exon 9 contained within the HDRT. HA-L/R: Homology-arm left/right; GSG: linker consisting of Gly-Ser-Gly; P2A: self-cleavage protein; pA: poly(A) sequence. (**B**) Integrase-mediated site-specific integration of the circular DNA donor vector into the GAPDH locus on DNA, mRNA and protein level. The promoter-less DNA donor with splice acceptor and P2A following the attV array (corresponding to the attG array on the HDRT) leads to RFP expression through the endogenous GAPDH promoter upon successful integration. Not drawn to scale.

**Suppl. Fig. 5.**
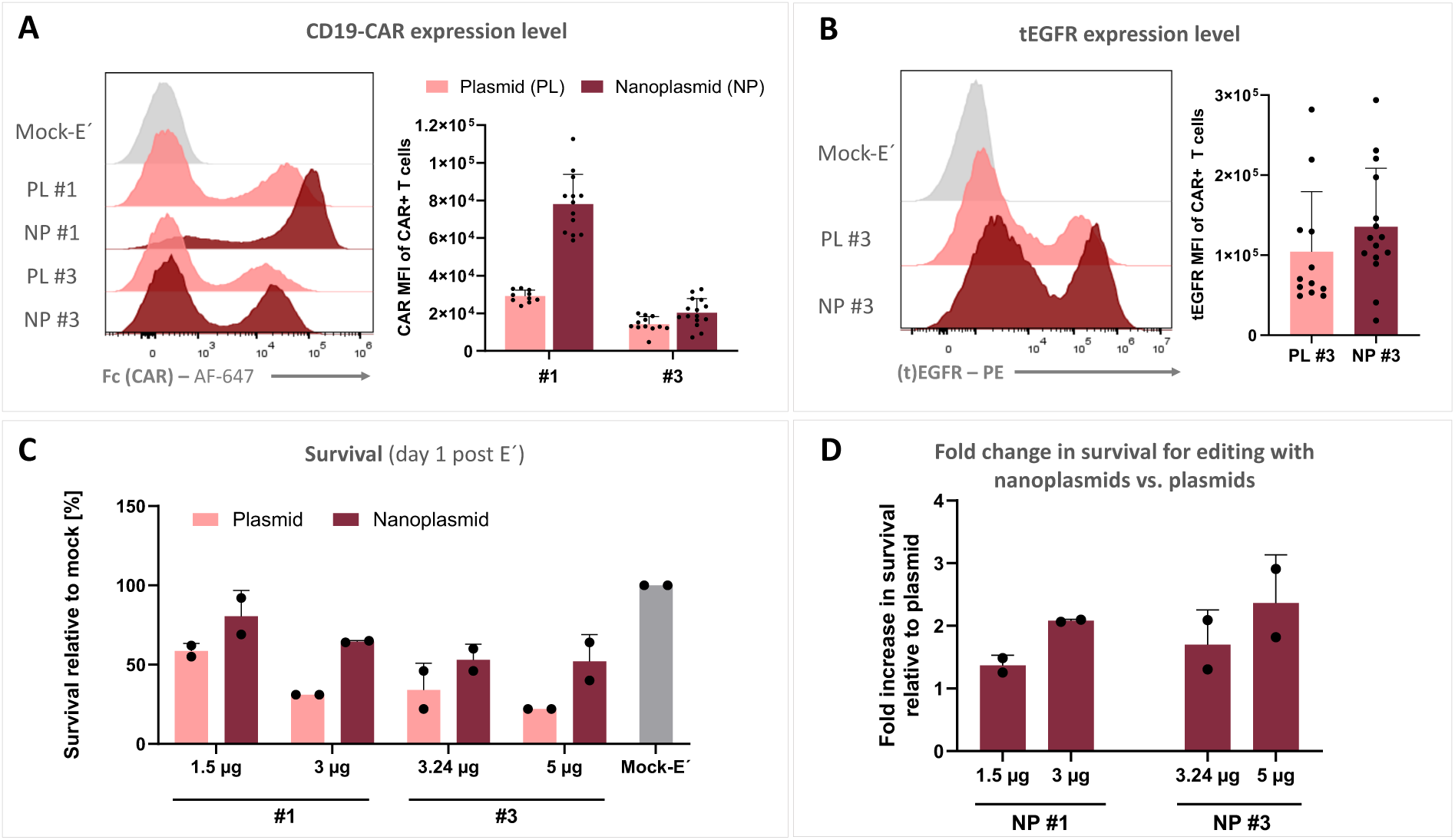
Nanoplasmids can improve transgene expression and cell viability of edited T cellst. (**A**) CD19-CAR expression levels for the monocistronic construct (#1) and tricistronic construct (#3) delivered as standard plasmid (PL) or nanoplasmid (NP). Median fluorescent intensity (MFI) of CD19-CAR expression in CAR+ T cells was determined by flow cytometry on day 4 after electroporation (n = 6 healthy donors). (**B**) Expression level of tEGFR in CAR+ T cells edited with PL #3 or NP #3 (n = 6 healthy donors). (**C)** Cell viability on day 1 after electroporation, normalized to mock-electroporation, measured in two healthy donors. (D) Fold increase in T cell viability following editing with nanoplasmids compared to standard plasmids (n = 2 healthy donors).

**Suppl. Fig. 6.**
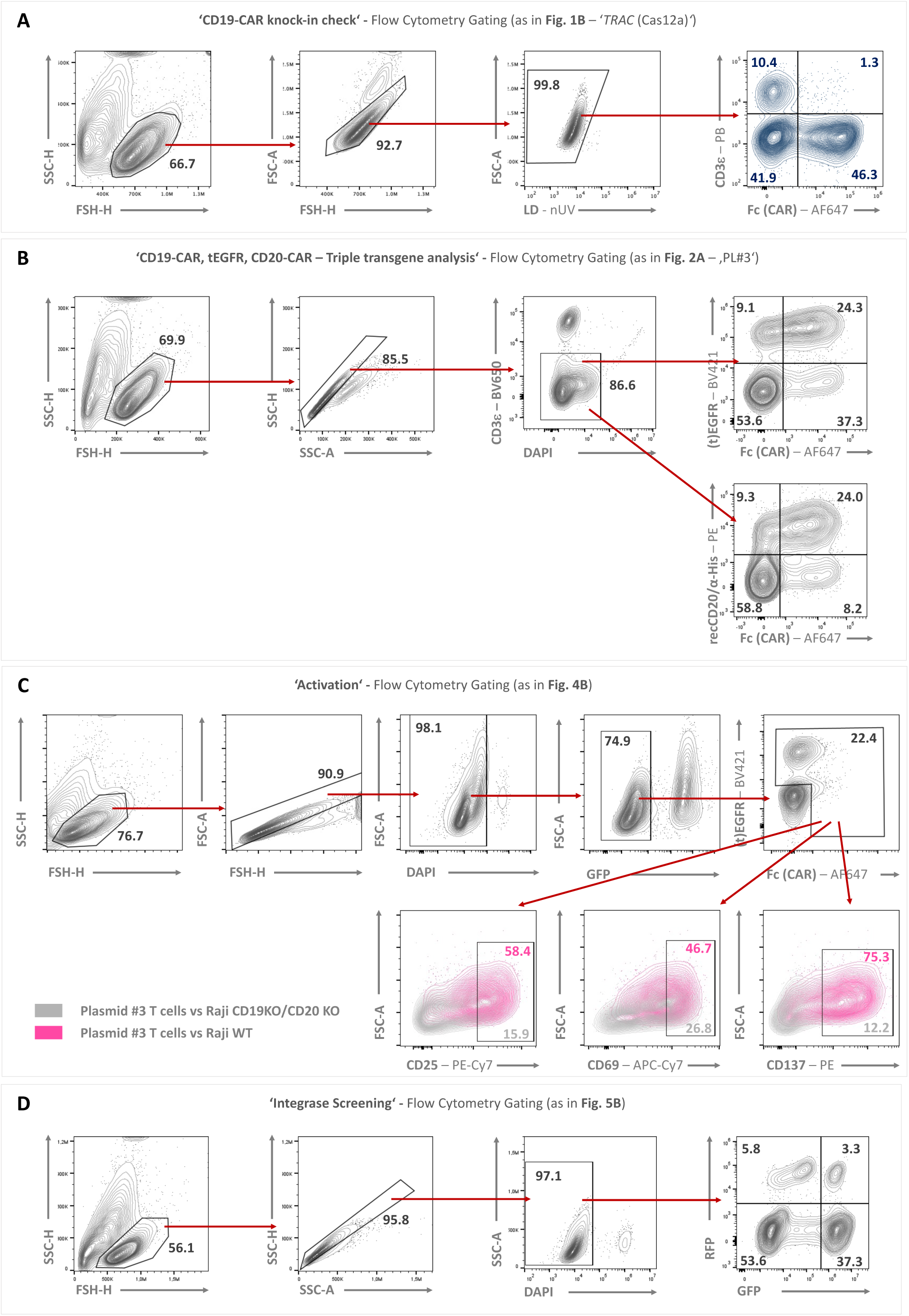
Gating Strategies.

